# A Streamlined Protocol for Small-scale Protoplast Generation and CRISPR/Cpf1-mediated Genome Editing in *Fusarium oxysporum*

**DOI:** 10.64898/2026.01.28.702186

**Authors:** Jun-Ze Zheng, Szu-Chieh Huang, Wen-Ting Zeng, Ying-Hong Lin, Tao-Ho Chang

**Author notes:** No. 15, Guangming Rd., Nantou City, Nantou County, 540, Taiwan (Tao-Ho Chang).

## Abstract

*Fusarium oxysporum* is a significant threat to agriculture and One Health, requiring advanced molecular tools for functional genomics analyses and biological control agent development. Existing gene-editing methods are hampered by costly protoplast preparation protocols and by CRISPR/Cas9 limitations, such as restricted PAM sequences and complex guide RNA requirements. We engineered an efficient CRISPR/Cpf1 system that overcomes these issues through three main innovations: small-scale protoplast generation using novel filter columns that greatly reduces enzyme consumption while simplifying workflows, a CRISPR/Cpf1 system with flexible PAM recognition and staggered DNA cleavage to promote homologous recombination, and minimal homology arm strategies that significantly decrease cloning complexity. Extensive validation confirms successful gene targeting with molecular verification and functional analysis via standardized pathogenicity assays. This integrated platform offers affordable, accessible tools for systematic *F. oxysporum* research, enhancing fundamental understanding of plant-pathogen interactions and supporting high-throughput screening vital for agricultural biotechnology and biological agent development.

**MOTIVATION:** *Fusarium oxysporum* plays crucial roles as both a damaging plant pathogen and a model for studying host-pathogen interactions. It is well established that systematic functional genomics approaches are vital for advancing agricultural biotechnology and developing biological agents. Creating efficient gene-editing systems can help elucidate virulence mechanisms and enable rapid production of modified strains for practical use. However, current transformation methods face major challenges, such as high enzyme costs, and CRISPR/Cas9 systems are limited by PAM sequence availability and by blunt-end DNA cleavage, which hampers homologous recombination. Furthermore, complex guide RNA scaffolds complicate large-scale functional studies with traditional methods. To address these challenges, we have developed a streamlined CRISPR/Cpf1 (Cas12a) platform that combines small-scale protoplast preparation with significantly reduced enzyme use, exploiting Cpf1’s unique features, such as flexible PAM recognition, staggered DNA cuts that promote recombination, improved target specificity, and simpler guide RNA design. This platform can also accelerate the development of biological agents and support high-throughput screening applications essential to the progress of agricultural biotechnology.

*HIGH LIGHTS:*

- Reduced enzyme costs by 95% through small-scale protoplast preparation
- CRISPR/Cpf1 system established for efficient gene editing in *Fusarium oxysporum*
- Streamlined workflow enables routine gene targeting and rapid mutant screening
- Complete workflow validated with EGFP-marked pathogenicity

## INTRODUCTION

*Fusarium oxysporum* represents one of the most economically important soilborne plant pathogens worldwide, causing vascular wilt diseases that result in substantial annual crop losses.^1,2^ This highly adaptable pathogen exhibits remarkable host specificity through specialised forms (*formae speciales*) that have co-evolved with specific plant families, making it an ideal model system for understanding plant-pathogen interactions.^3^ Advancing knowledge of *F. oxysporum* virulence mechanisms and developing effective disease management strategies requires robust molecular tools for systematic gene function analysis, which are essential for both fundamental research and practical applications in crop protection and biotechnology.

Despite the crucial role of molecular research in *F. oxysporum*, progress in functional genomics faces major challenges due to technical barriers. Fungal transformation remains difficult, involving labour-intensive protoplast preparation with costly enzymatic mixes and precise execution.^4^ Conventional methods demand large amounts of cell wall-degrading enzymes, making large-scale experiments cost-prohibitive.^5^ Additionally, these workflows are time-consuming, often taking extended periods from protoplast creation to mutant confirmation. These challenges restrict the scope of systematic studies of gene function and slow progress in understanding the molecular basis of pathogenicity and host specificity.

Current CRISPR/Cas9 technology has significantly advanced fungal molecular biology, providing enhanced precision for targeted gene disruption and modification across diverse fungal species.^6^ CRISPR/Cas9 systems have been successfully implemented in several Fusarium species, enabling targeted gene knockouts and facilitating investigations of virulence mechanisms.^7–9^ However, traditional CRISPR/Cas9 approaches in fungi require extensive homology arms (typically 500-2000 bp) for reliable homology-directed repair.^10^ The blunt-end DNA cuts generated by Cas9 preferentially activate non-homologous end joining (NHEJ) repair pathways, which are error-prone and often result in imprecise gene modifications rather than targeted homologous recombination.^11^ These characteristics necessitate complex experimental designs and may limit the efficiency of precise gene targeting applications in fungal systems.

The emergence of CRISPR/Cpf1 (Cas12a) technology offers distinct advantages that may address these limitations.^12–14^ Cpf1 generates staggered DNA breaks with 4-5 nucleotide overhangs that promote homologous recombination and reduce reliance on error-prone NHEJ pathways. Additionally, Cpf1 requires only a simple TTTV protospacer adjacent motif compared to the restrictive NGG requirement of Cas9, expanding target site accessibility within fungal genomes.^14^ The system also utilises shorter guide RNAs that require no complex scaffold, potentially simplifying experimental protocols.^14^ Despite these theoretical advantages, the application of the CRISPR/Cpf1 system has not yet been established in *F. oxysporum*.

This study aims to develop and validate an optimised CRISPR/Cpf1 system for *F. oxysporum* transformation that addresses current technical and economic limitations. Our objectives were to establish streamlined protoplast preparation protocols, systematically evaluate Cpf1-mediated gene editing capabilities, and integrate these innovations into a comprehensive workflow suitable for high-throughput functional genomics applications.

## RESULTS

### Filter Column-based Design for Small-scale Protoplast Isolation

This novel filter column-based approach streamlines the entire protoplast preparation workflow and significantly reduces enzyme consumption, addressing the complexity and cost barriers of traditional methods (Scheme 1). The design employs a disposable filter column as both reaction vessel and separation device, allowing gentle processing of fungal mycelium through controlled washing, digestion, and collection steps. The protocol begins with loading overnight-grown Fusarium mycelium directly into the filter column, followed by sequential washing to remove growth medium and spores. Enzymatic digestion occurs within the inverted column under gentle rotary shaking, and protoplasts are recovered through controlled centrifugation steps that minimize mechanical stress while maximizing yield.

**Scheme 1.**
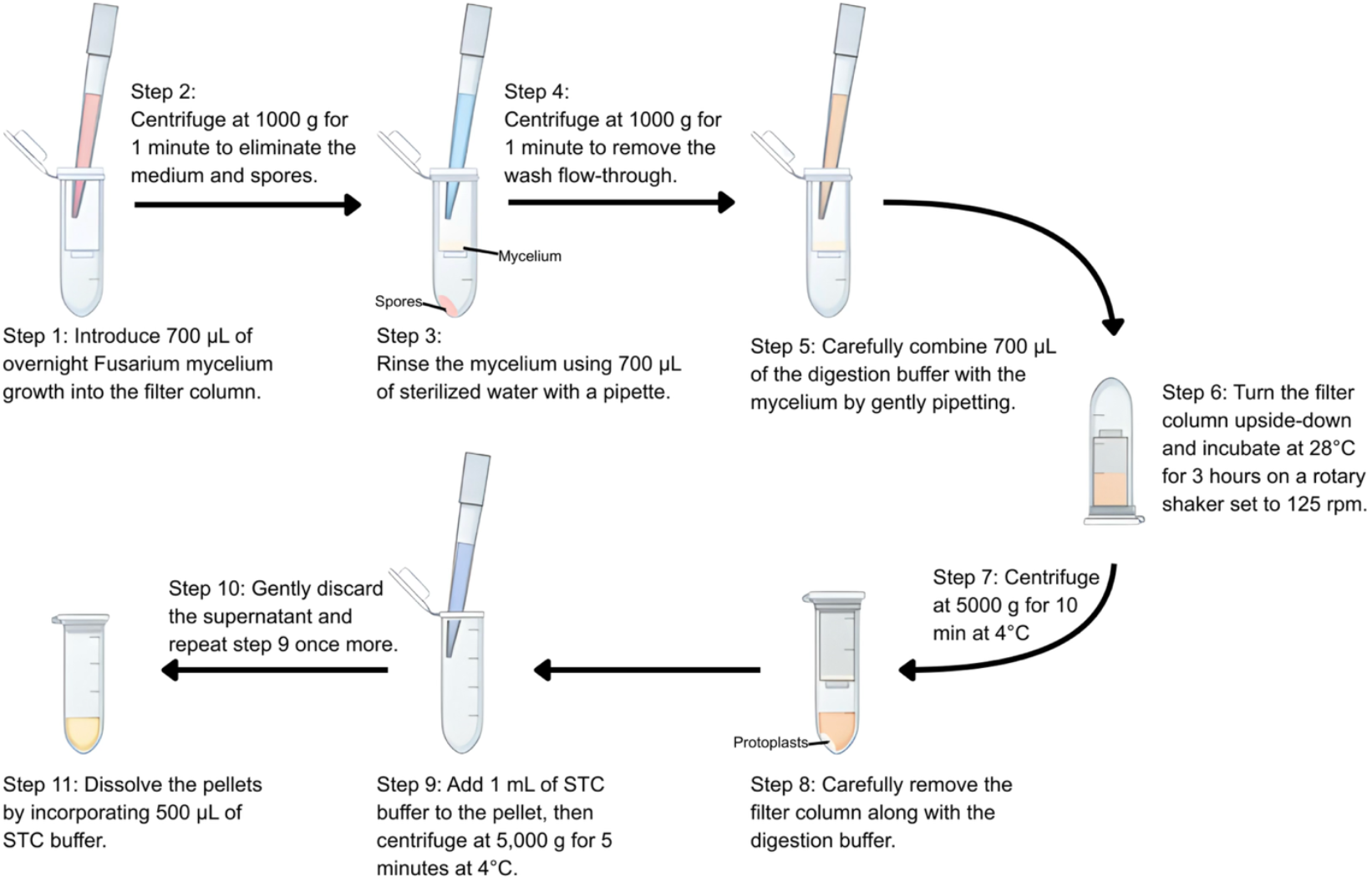
Filter column-based design for small-scale protoplast isolation. Step-by-step workflow showing mycelium loading into filter columns, washing procedures, enzymatic digestion in inverted columns with rotary shaking, centrifugation-based protoplast recovery, and final resuspension in SuTC buffer.

Protoplast isolation optimisation revealed significant performance differences across experimental conditions (Figure 1 and Figure S1). Buffer system comparison demonstrated that both 0.8 M NaCl and 0.8 M KCl solutions yielded comparable protoplast densities (median ∼25-30 × 10^6^ cells/mL), significantly outperforming phosphate-based and mannitol-based buffers, which achieved less than 5 × 10^5^ cells/mL (Figure 2A). Systematic enzyme concentration analysis showed that the filter column approach achieved optimal protoplast production (highest yield around 6.0 × 10^7^ cells/mL at 2 hours) with 7 mg Driselase and 10 mg β-glucanase per 0.7 mL reaction (Figure 2B).

**Figure 1.**
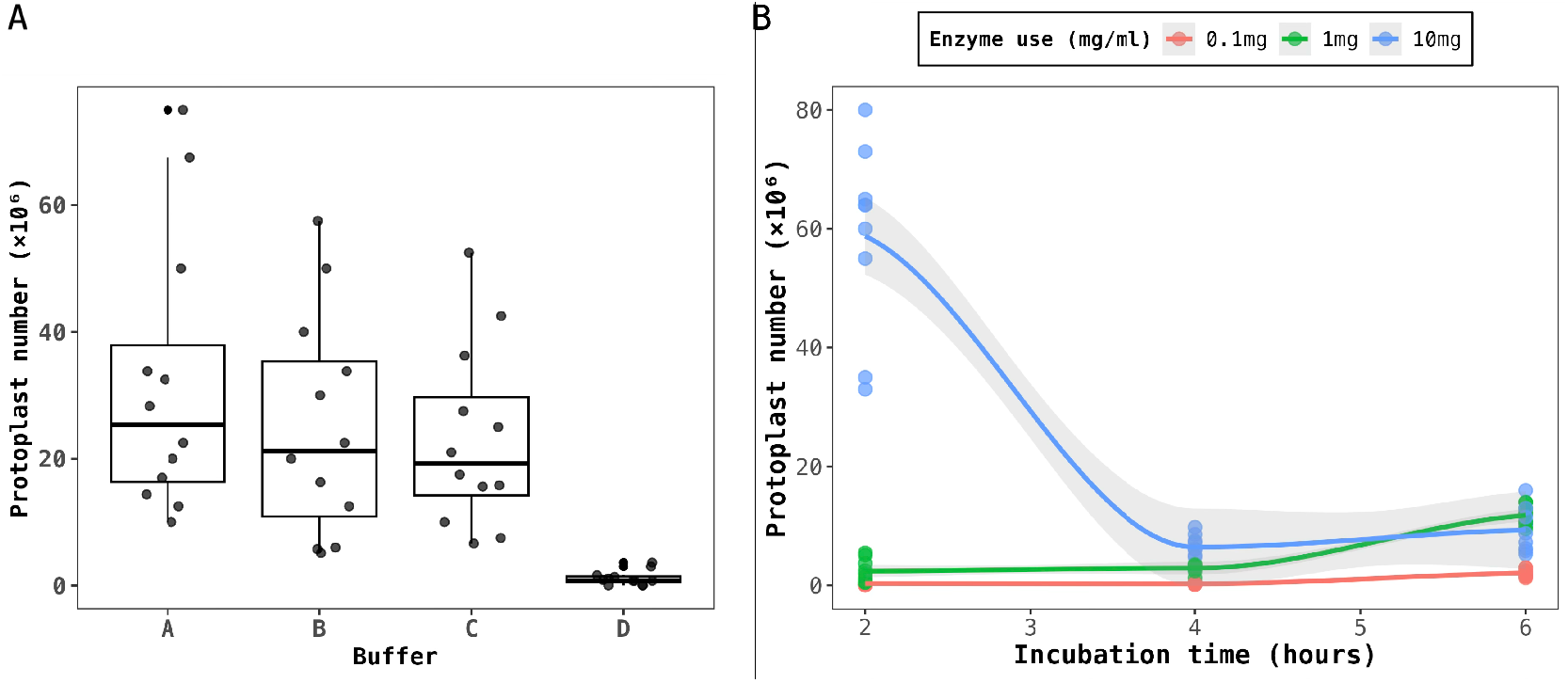
Protoplast isolation experiments. (A) Protoplast yields using different buffer systems: A, 0.8 M NaCl; B, 0.8 M KCl; C, 10 mM Na_2_HPO_4_, 20 mM CaCl_2_, 1.2 M NaCl, pH 5.8; D, 0.6 M mannitol, 10 mM Tris-HCl, 10 mM CaCl_2_, pH 7.5. Box plots show median, quartiles, and individual data points. (B) Time-course analysis of protoplast production using different Driselase concentrations (0.1 mg, 1 mg, and 10 mg per reaction) over 2-6 hours incubation. Points represent individual measurements with trend lines and confidence intervals.

**Figure 2.**
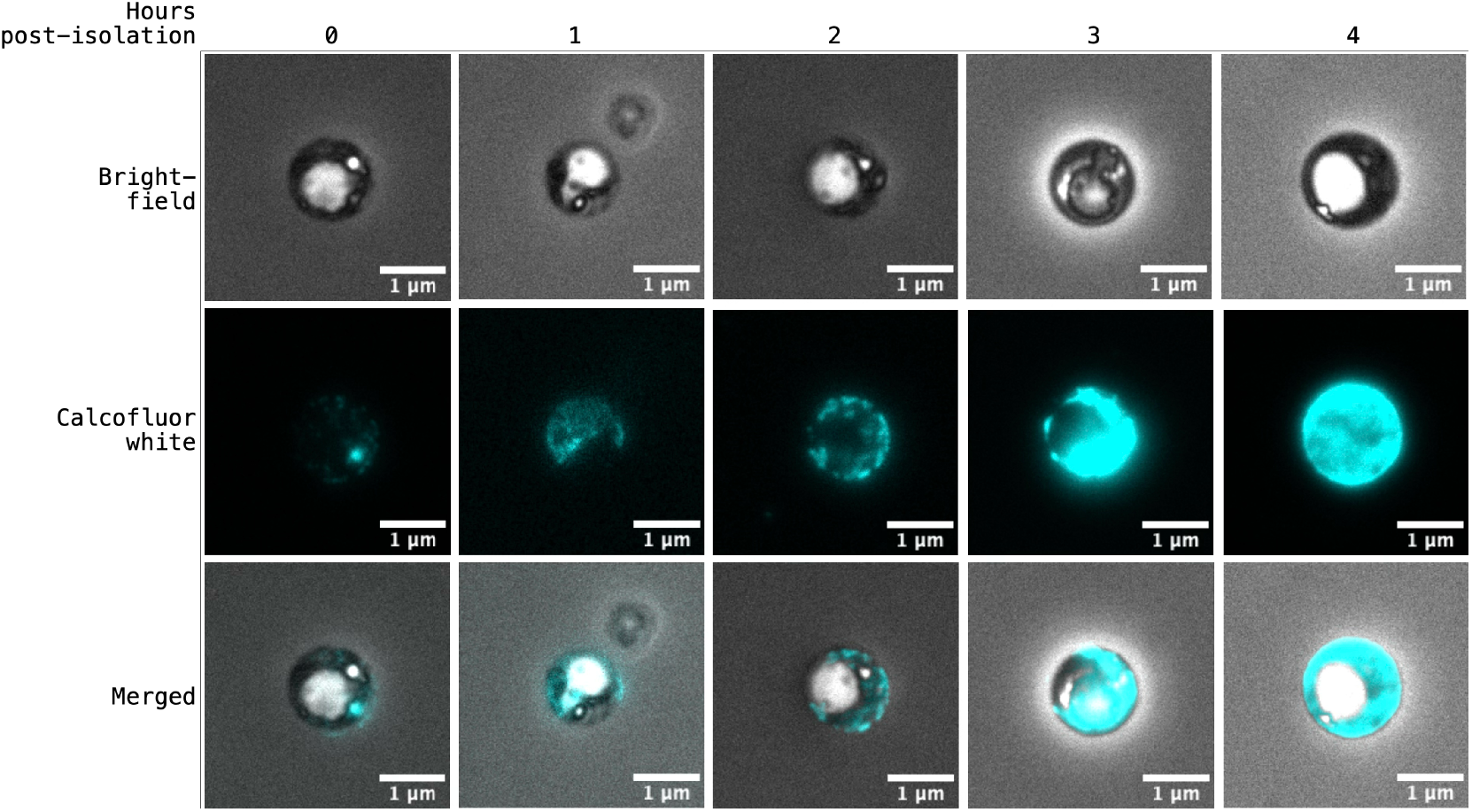
Time-course analysis of cell wall regeneration. Representative protoplasts showing cell wall reformation at 0, 1, 2, 3, and 4 hours post-isolation. Images captured under bright-field illumination (top row), calcofluor white fluorescence staining (middle row), and merged channels (bottom row). Calcofluor white specifically stains cellulose and chitin components of fungal cell walls. Scale bars, 1 μm.

Time-course analysis of cell wall regeneration showed quick rebuilding patterns essential for transformation timing (Figure 2). Calcofluor white staining indicated that freshly isolated protoplasts had minimal cell wall material at 0 hours post-isolation, with gradual wall rebuilding visible by 1-2 hours and complete reformation by 4 hours. This swift regeneration process requires immediate transformation after protoplast preparation to achieve maximum efficiency.

### Systematic Design and In Vitro Validation of URA5-targeting gRNAs for CRISPR/Cpf1

The *URA5* gene was selected as the CRISPR/Cpf1 target due to its well-characterised phenotype and utility as a negative selection marker (Figure 3). gRNA design and validation identified two optimal target sites within the URA5 coding sequence, with both gRNA_244657 and gRNA_245247 demonstrating robust in vitro cleavage activity. Concentration-dependent analysis established optimal working concentrations between 1-3 μM, with detectable activity maintained at concentrations as low as 0.15 μM.

**Figure 3.**
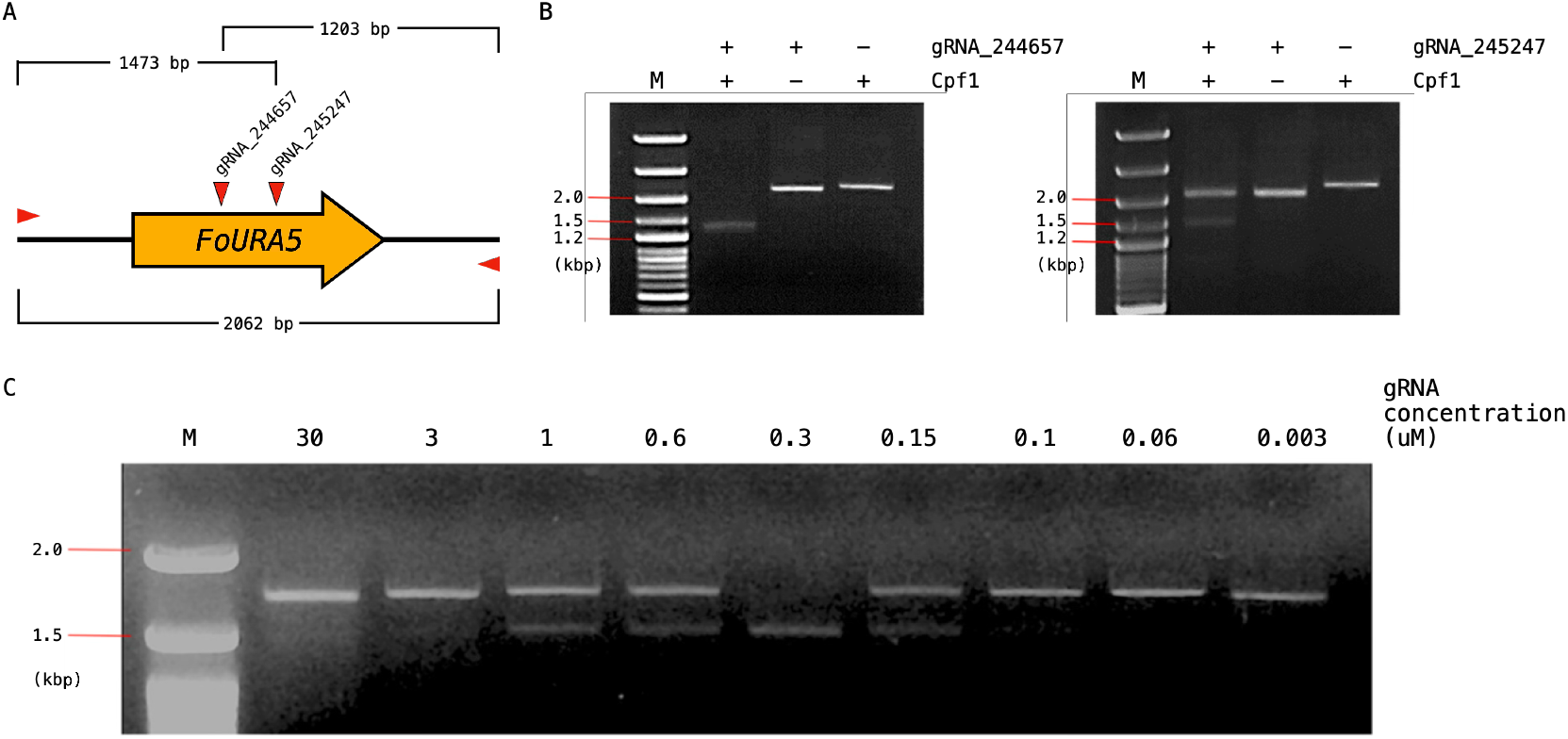
Systematic design and *in vitro* validation of *URA5*-targeting gRNAs for CRISPR/Cpf1. (A) Schematic diagram illustrating *the FoURA5* gene structure, with target sites for gRNA_244657 and gRNA_245247 marked by arrows. The expected cleavage fragments are 1203 bp for gRNA_244657 and 1473 bp for gRNA_245247, with the total genomic region spanning 2062 bp. Primer binding sites are indicated with red triangles. (B) *In vitro* cleavage assays for both gRNAs using purified genomic DNA. Reactions were performed with (+) and without (-) gRNA or Cpf1 components. M denotes a 1 kbp DNA ladder. (C) Concentration-dependent cleavage analysis demonstrating gRNA activity across serial dilutions from 30 μM to 0.003 μM. M represents a 100 bp DNA ladder.

### GFP-based Screening Reveals Homology Arm Length-dependent Editing Efficiency

Systematic evaluation of homology arm requirements using EGFP reporter integration revealed clear differences in efficiency across arm lengths (Figure 4 and Figure S). Transformants with 1000-bp homology arms exhibited 87.4% EGFP-positive colonies, whilst 100-bp arms achieved 69.8% efficiency. Most significantly, minimal 20-bp homology arms supported 49.5% EGFP integration, demonstrating that extremely short homology sequences retain functional gene targeting capability.

**Figure 4.**
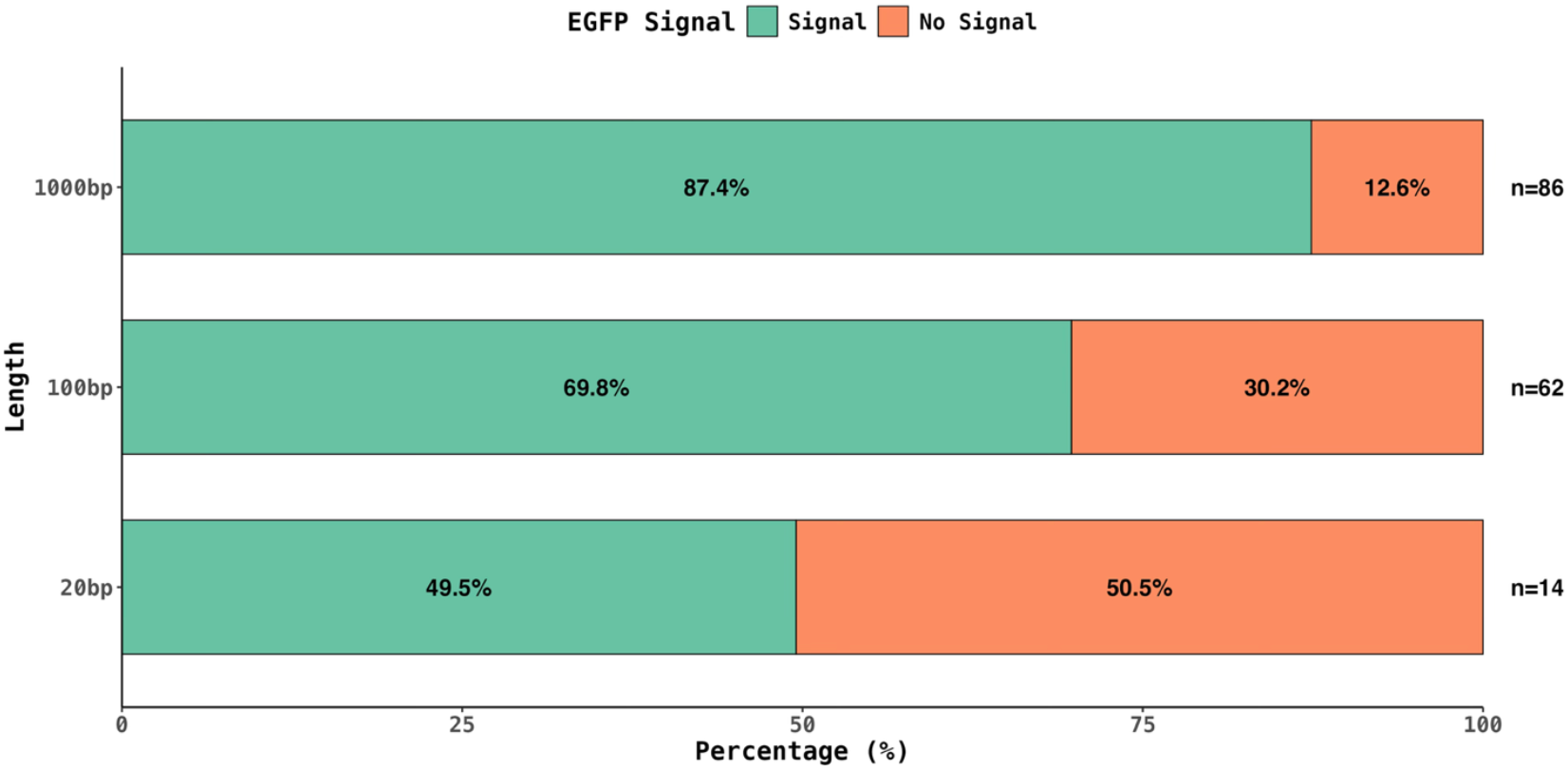
Screening for *ΔURA5* Lines with EGFP signals using different homology arm lengths. Proportion of EGFP-positive colonies among 5’-FOA resistant transformants using donor constructs with different homology arm lengths (20 bp, 100 bp, and 1000 bp). All colonies were pre-selected on 5’-FOA medium, indicating successful *URA5* gene disruption and uracil auxotroph. Stacked horizontal bars show percentage of ***ΔURA5*** colonies with EGFP signal (green, indicating donor integration) and without EGFP signal (orange, indicating URA5 disruption without donor integration). Sample sizes (n) indicated for each condition.

### Molecular Confirmation Reveals Precise Editing Event Classification

A comprehensive genotyping analysis was performed using a three-primer-pair system to systematically classify editing outcomes (Figure 5). The primer design allowed differentiation between precise integration, partial insertions, and indel formation (Figure 5A). Classification of mutation events showed that using 20-bp homology arms resulted in 13.1% of all transformants with accurate simple insertions, while longer homology arms increased the accuracy proportionally (Figures 5B,C). A comparative analysis with hygromycin resistance selection revealed mostly random integration events (Figure S3), highlighting the importance of the selection method for reliable HDR detection.

**Figure 5.**
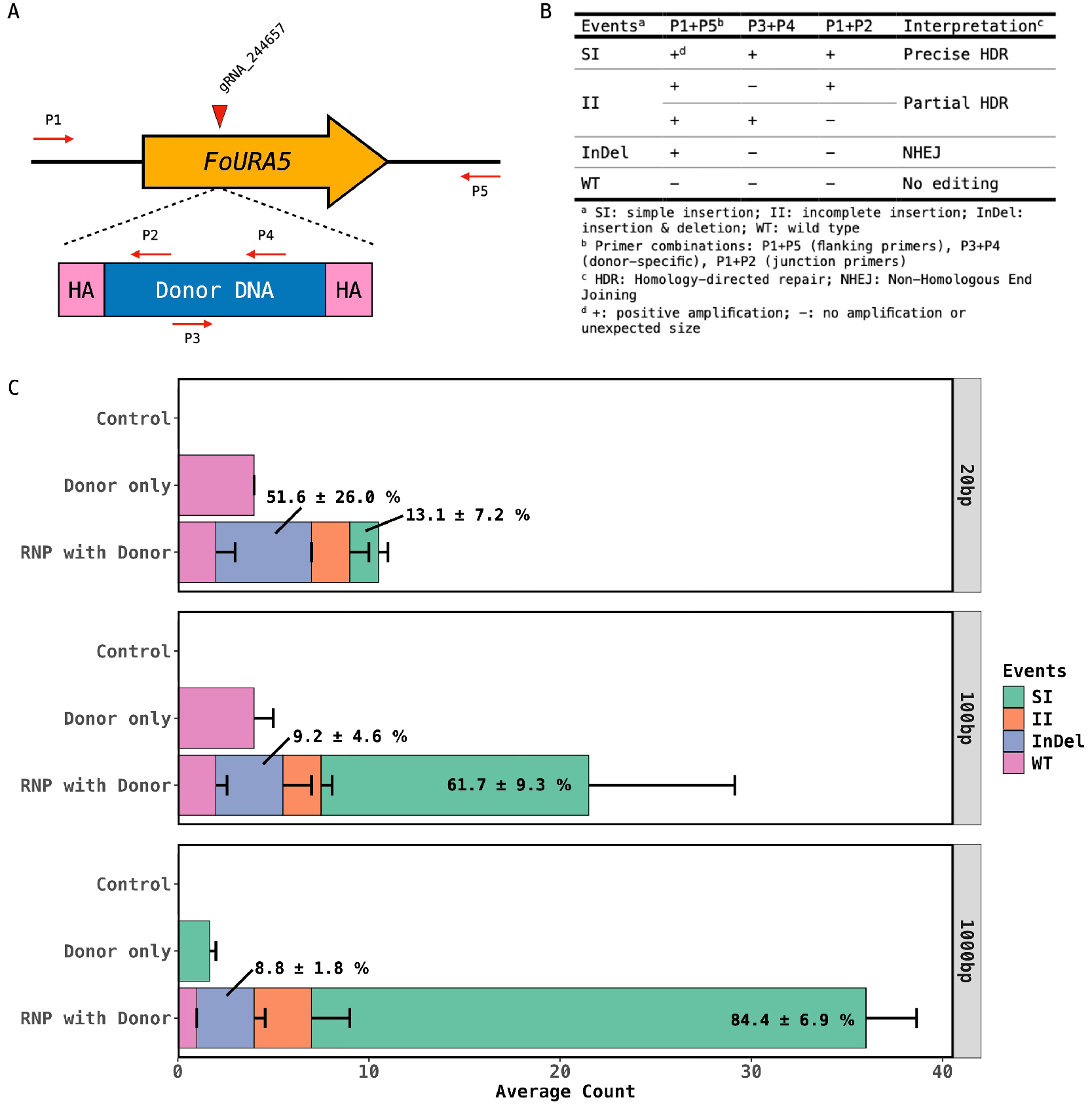
Genotyping for *URA5* editing events. (A) Primer design scheme for genotyping analysis. Five primers (P1-P5) are positioned to identify different editing outcomes through PCR amplification patterns. P1+P5 detect gene disruption, P3+P4 identify donor DNA presence, and P1+P2 verify homologous recombination junctions. (B) Classification table for editing events based on PCR results from three primer combinations. SI, simple insertion; II, incomplete insertion; InDel, insertion and deletion; WT, wild type. (C) Quantitative analysis of editing outcomes with different homology arm lengths (20 bp, 100 bp, 1000 bp). Bars display average counts of each editing event type among transformants under three conditions: Control, Donor only, and RNP with Donor. Error bars indicate standard error of three repeat experiments. Colour coding corresponds to different mutation types as defined in panel B.

### Validation Platform Demonstrates Pathogenicity Assessment Capability

Standardised inoculation assays verified the *Fusarium* pathogenicity platform’s ability to distinguish pathogen specificity (Figure 6). FCN33 (*F. oxysporum* f. sp. *conglutinans*, crucifer-adapted) caused severe symptoms like yellowing and stunted growth, while Foc24 (*F*. oxysporum f. sp. cubans, banana-adapted) demonstrated pathogenicity on Arabidopsis. This confirms the platform can effectively identify host-specific differences.

**Figure 6.**
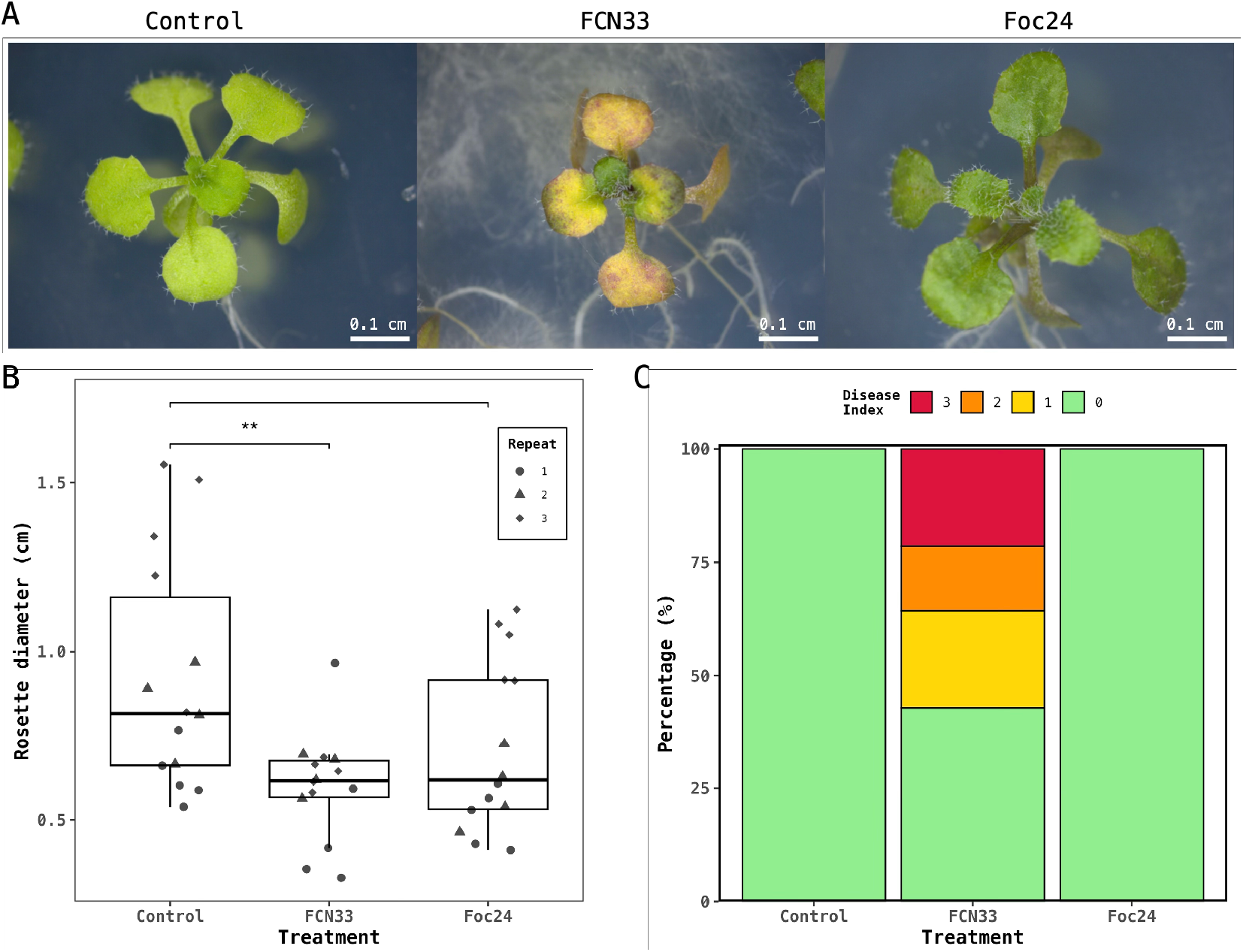
Arabidopsis inoculation platform for host specificity assessment. (A) Representative *Arabidopsis thaliana* seedlings at 7 days post-inoculation showing disease symptom development. Control, mock-inoculated plants; FCN33, crucifer-adapted *F. oxysporum* f. sp. *conglutinans*; Foc24, banana-adapted *F. oxysporum* f. sp. *cubense*. Scale bars, 0.1 cm. (B) Quantitative analysis of rosette diameter across treatment groups. Box plots show median, quartiles, and individual data points. Sta (C) Disease index showing percentage distribution of symptom categories across treatments. The disease index as follows: 0: less than 30% wilt leaves; 1: 30%∼50% wilt leaves; 2: 50%∼70%; 3 over 70% wilt leaves.

Microscopic analysis of edited strain colonisation demonstrated successful root invasion patterns (Figure 7). EGFP-expressing *URA5* edited strains exhibited extensive hyphal networks within Arabidopsis root tissues, confirming that genetic modification did not impair infection capability whilst providing real-time visualisation of colonisation processes.

**Figure 7.**
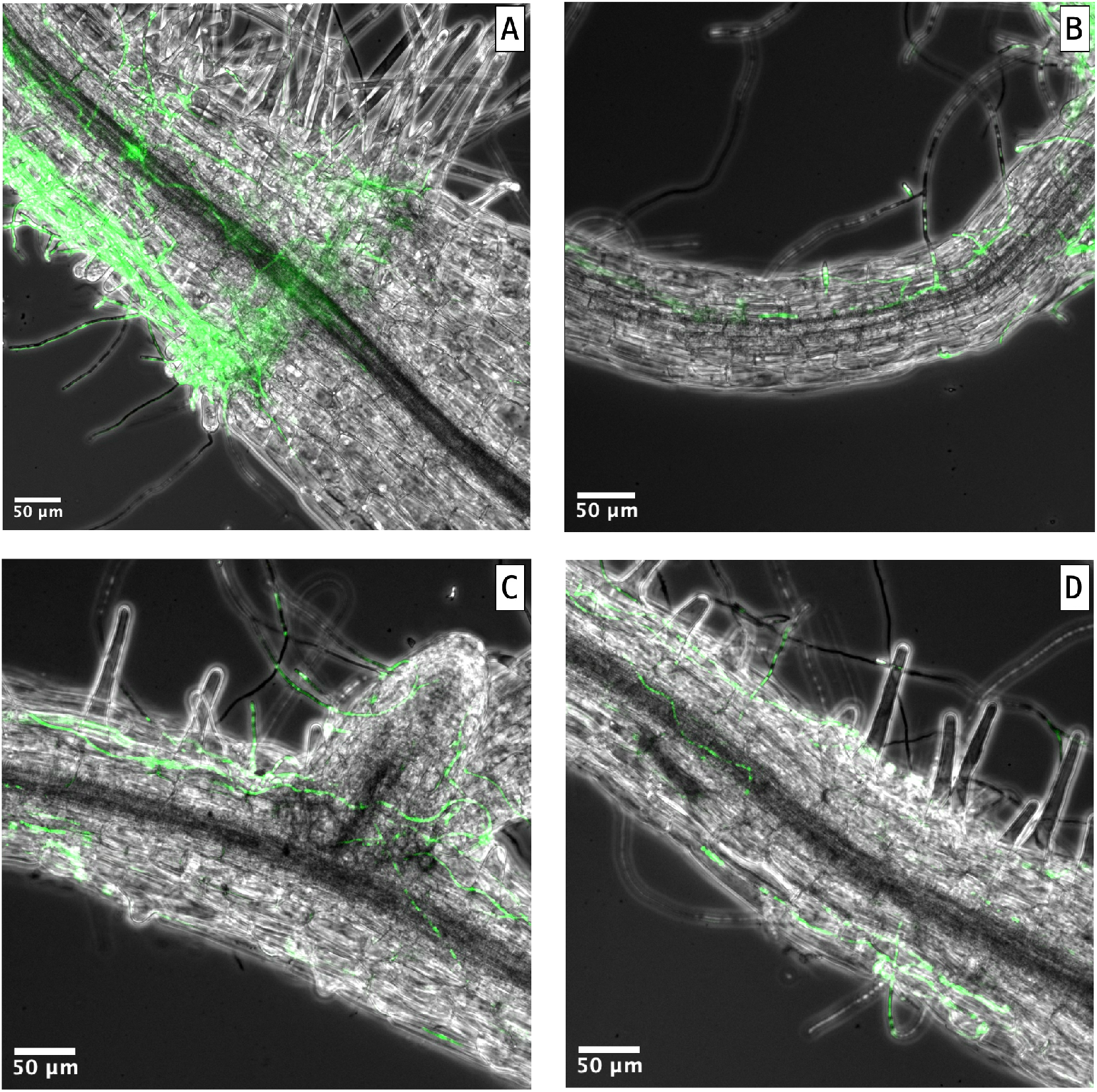
Fluorescence microscopic examination of *URA5*-edited strain in Arabidopsis root colonisation. (A-D) Fluorescence microscopy images show EGFP-expressing URA5-edited *F. oxysporum* strain colonising *Arabidopsis thaliana* roots. Green indicates fungal hyphae; grey shows plant tissue under bright-field. (A) Hyphal clustering near shoot base. (B) Hyphal growth between plant cells. (C) Colonisation at lateral root junctions. (D) Hyphal penetration within vascular tissue. Scale bars are 50 μm.

## DISCUSSION

The small-scale protoplast preparation notably reduces enzyme consumption while maintaining comparable yields, requiring only 7 mg of Driselase compared to 70-800 mg in traditional protocols across various *Fusarium* species (Table 1). This significant reduction in enzyme use, combined with simplified small-scale processing, overcomes a major economic barrier that has limited large-scale functional studies.

**Table 1.** Comparison of Driselase usage in fungal protoplasts preparation.

| Study | Organism/Strain | Driselase (mg) | Volume (mL) | Yield (x10<sup>8</sup>/mL) | Additional enzymes |
| --- | --- | --- | --- | --- | --- |
| This study | *Fusarium oxysporum* f. sp. *conglutinans* | 7 | 0.7 | 0.3-0.8 | β-glucanase |
| Yang et al. (2025)<sup>15</sup> | *F. oxysporum* f. sp. *fragariae* (H6) | 70-100 | 10 | Not reported | Cellulase, lysozyme |
| Shi et al. (2025)<sup>16</sup> | *F. oxysporum* (FO-1) | 500 | 25 | 1.44 | none |
| Ayukawa et al. (2021)<sup>17</sup> | *F. oxysporum* f. sp. *conglutinans* | 500 | 25 | 4 | Lysing enzymes |
| Shao et al. (2024)<sup>18</sup> | *F. graminearum* (PH-1) | 100 | 20 | Not reported | Lysing enzymes, snailase |
| Hicks et al. (2023)<sup>19</sup> | *F. graminearum* (DAOMC233423) | 500 | 20 | 2 | Lysing enzymes, yatalase |
| Lohmar et al. (2022)<sup>20</sup> | *F. verticillioides* (Fv999) | 800 | 20 | 5-10 | Lysing enzymes, chitinase |

The faster workflow, enabling edited strains to be obtained within days instead of weeks, results from integrated rapid monitoring of cell wall regeneration and more efficient transformation protocols. These combined enhancements offer considerable advantages for systematic functional studies, potentially allowing researchers to perform gene function analyses that were previously restricted by resource constraints.

Our demonstration that 20-bp homology arms support functional gene targeting represents a significant advancement over conventional CRISPR approaches, which typically require 500-2000 bp homology arms for reliable HDR in fungi.^21^ This capability stems from key biochemical differences between Cpf1 and Cas9 systems. Most significantly, Cpf1 generates staggered DNA breaks with 4-5 nucleotide overhangs that facilitate homologous recombination, whereas Cas9 produces blunt-end cuts that preferentially activate error-prone NHEJ pathways.^22^ Additionally, Cpf1’s TTTV PAM requirement offers greater targeting flexibility compared to Cas9’s restrictive NGG motif, whilst utilising shorter guide RNAs without complex scaffold requirements.^23^

The 13.1% precise HDR efficiency achieved through our URA5/5’-FOA selection strategy demonstrates that these advantages translate into practical benefits, with the two-step selection approach effectively enriching for editing events. Nevertheless, the utilisation of minimal homology arms also increases the likelihood of random integration events. Our analysis revealed that although 20-bp arms facilitate functional targeting, they may also promote non-specific insertion, particularly with intense selection systems (i.e. Hyg^R^ system) (Figure S3). Traditional preparation of lengthy homology arms requires several weeks of cloning, whereas 20-bp sequences can be incorporated via oligonucleotide synthesis within hours,^24^ thereby significantly accelerating experimental timelines. The trade-off between cloning simplicity and integration precision underscores the importance of optimising the selection system, as an initial analysis with Hygromycin selection revealed extensive random insertion events compared with the more selective URA5/5-FOA approach.

The platform’s utility extends beyond fundamental research to support agricultural biotechnology objectives. The generation of an EGFP-marked strain demonstrates real-time monitoring capabilities for host-pathogen interaction studies.^25^ This enables dynamic visualisation of colonisation processes that advance traditional plant pathology methodologies. The established Arabidopsis inoculation system provides a standardised phenotypic validation framework, facilitating the assessment of gene function and spatial expression patterns during pathogenesis.^26^ The combination of rapid mutant generation, cost-effective screening, and reliable validation creates a framework for systematic knockout studies that provides a foundation for identifying virulence factors and potential biocontrol targets, supporting biotechnology development tailored to agricultural challenges.

### Limitations of the study

Several constraints limit the broader applicability of this platform. Protoplast-based transformation remains technically demanding due to cell fragility and regeneration variability, impacting reproducibility particularly in high-throughput applications. The 13.1% precise HDR efficiency with 20-bp arms, whilst functional, may limit applications requiring high-frequency modification. A significant limitation is the dependence on selection system design: favourable results with URA5 benefit from 5-FOA negative selection, but targeting other genes would rely primarily on EGFP screening, potentially requiring screening of significantly more transformants to identify precise editing events. Current validation limited to URA5 in *F. oxysporum* necessitates comprehensive evaluation across diverse targets and fungal species to establish general utility.

## RESOURCE AVAILABILITY

### Lead contact

Requests for further information and resources should be directed to and will be fulfilled by the lead contact, Tao-Ho Chang.

### Materials availability

- This study did not generate new unique reagents.
- All unique/stable reagents generated in this study are available from the lead contact without restriction.

### Data and code availability

- All data reported in this paper will be shared by the lead contact upon request.
- This paper does not report original code.
- Any additional information required to reanalyze the data reported in this paper is available from the lead contact upon request.

## ACKNOWLEDGMENTS

This work was financially supported by the National Science and Technology Council (NSTC), Taiwan, under grant numbers 112-2313-B-005-008-, 114-2313-B-005-018-MY3 and 114-2637-B-020-002-. We extend sincere appreciation to colleagues at the Academy of Circular Economy, National Chung Hsing University, for their invaluable guidance and technical assistance throughout this research. Special thanks are given to Prof. Jenn-Wen Huang and Prof. Pi-Fang Linda Chang for their essential contributions during the initial stages of this study, including the provision of *F. oxysporum* strains and critical research materials that enabled this investigation. Additional acknowledgement is given to the technical staff and laboratory personnel who provided experimental support and assistance with laboratory procedures.

## AUTHOR CONTRIBUTIONS

Conceptualisation, J.-Z. Zheng and T.-H. Chang; methodology, A.B., S.C.P., and S.Y.W.; Investigation, J.-Z. Zheng, S.-C. Huang and W.-T. Tseng; writing—original draft, J.-Z. Zheng; writing—review & editing, Y.-H. Lin and T.-H. Chang; funding acquisition, Y.-H. Lin and T.-H. Chang; resources, Y.-H. Lin and T.-H. Chang; supervision, T.-H. Chang.

## DECLARATION OF INTERESTS

The authors declare that they have no competing interests.

## DECLARATION OF GENERATIVE AI AND AI-ASSISTED TECHNOLOGIES

During the preparation of this work, the author(s) used Claude (sonnet 4.5) to improve readability. After using this tool or service, the author(s) reviewed and edited the content as needed and take(s) full responsibility for the content of the publication.

## SUPPLEMENTAL INFORMATION

**Document S1. Figures S1–S3, Tables S1 and S2, and supplemental references**

## STAR★METHODS

### KEY RESOURCES TABLE

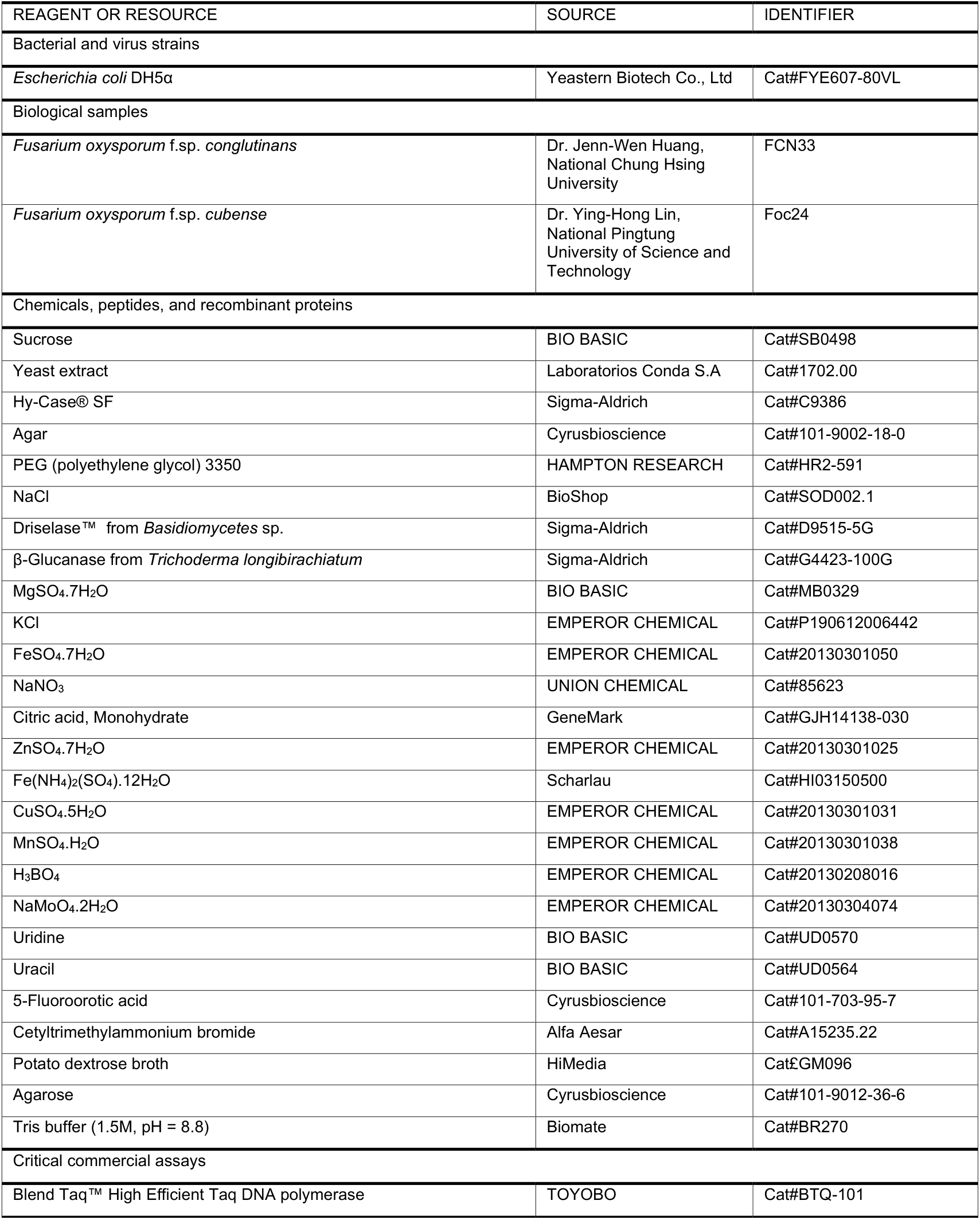
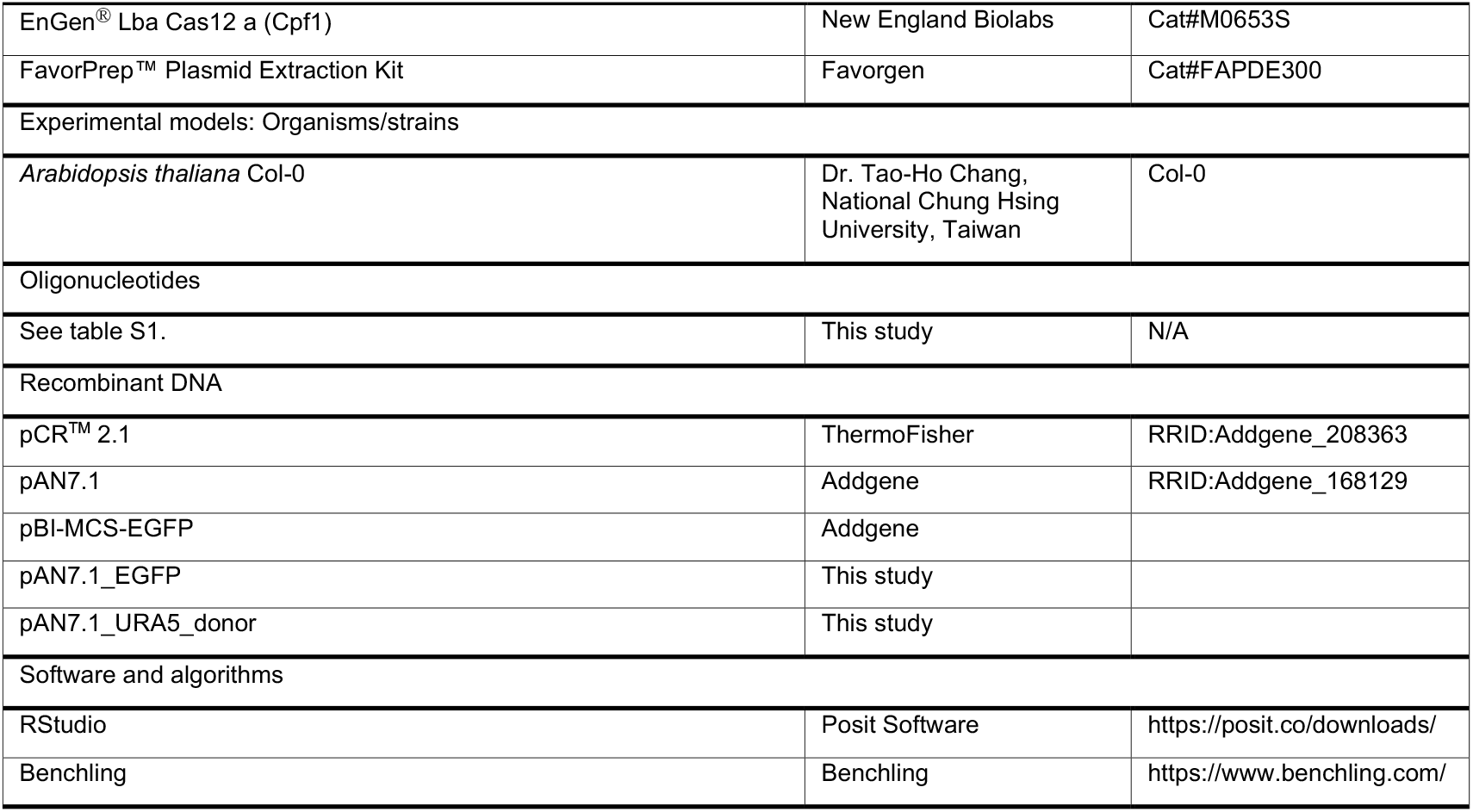

## METHOD DETAILS

### Experimental organisms and culture conditions

#### Fungal Strains

*Fusarium oxysporum* f. sp. *conglutinans* FCN33 (crucifer-adapted) and *F. oxysporum* f. sp. *cubense* race 4 Foc24 (banana-adapted) were used in this study. FCN33 was obtained from Dr. Jenn-Wen Huang (National Chung Hsing University, Taiwan) and Foc24 was provided by Dr. Ying-Hong Lin (National Pingtung University of Science and Technology). Single spore isolates were obtained using sterile glass needles under a dissecting microscope to ensure genetic homogeneity and culture purity. Stock cultures were preserved in sterile sand tubes containing 2% water agar and approximately 1% sandy soil, stored at room temperature for long-term maintenance. For routine culturing, strains were grown on half-strength potato dextrose agar (1/2 PDA: 19.5 g PDA per litre) at 28°C.

#### Plant Material

*Arabidopsis thaliana* Col-0 seeds were surface-sterilised for the plate assay according to the published protocols.^27^ Seeds were stratified at 4°C for 24 hours before sowing on half-strength Murashige and Skoog (MS) medium containing 1% agar and 1% sucrose. Plants were grown in a controlled environment chamber at 25°C under 16-hour photoperiod.

#### Culture Media

M100 selective medium (3% w/v sucrose, 0.5 g/L MgSO_4_.7H_2_O, 0.5 g/L KCl, 10 mg/L FeSO_4_.7H_2_O, 2 g/L NaNO_3_, 2 mL/L trace elements, 2% w/v agar) supplemented with appropriate selection agents. For URA5 selection, 2 mg/mL 5’-fluoroorotic acid (5’-FOA) was added along with 250 μM uridine and uracil. SuTC buffer (20% w/v sucrose, 50 mM Tris-HCl pH 8.0, 50 mM CaCl2·H_2_O) was used for protoplast manipulation and storage.

### Small-scale protoplast preparation

Mycelium preparation for protoplast isolation was initiated by inoculating 10^7^ conidia in 50 mL half-strength PDB and incubating at 28°C with 125 rpm shaking for 16-20 hours to obtain young mycelium.

The culture was filtered through a spin filter column (FavorPrep Genomic DNA Extraction Mini Kit, FAVORGEN) at 1,000 × g for 30 seconds to remove growth medium and residual conidia, followed by washing with 700 μL sterile distilled water.

Enzymatic digestion buffer preparation involved testing four different osmotic stabilising solutions: (A) 0.8 M NaCl, (B) 0.8 M KCl, (C) 10 mM Na_2_HPO_4_, 20 mM CaCl_2_, 1.2 M NaCl, pH 5.8, and (D) 0.6 M mannitol, 10 mM Tris-HCl, 10 mM CaCl_2_, pH 7.5. Each digestion buffer contained 10 mg Driselase (Sigma-Aldrich D9515) and 15 mg β-glucanase (Sigma-Aldrich G4423) per 1 mL total volume. All digestion buffers were filter-sterilised through 0.22 μm disc filters before use.

Protoplasts were generated by adding 700 μL of digestion buffer to the washed mycelium in the filter column. The mixture was gently pipetted until mycelium was fully suspended, then the column was inverted and incubated at 28°C with 125 rpm rotary shaking for 2 hours. Following digestion, the column was centrifuged at 5,000 × g for 10 minutes at 4°C to collect protoplasts in the collection tube. The filter column was carefully removed and the protoplast pellet was washed twice with 1 mL SuTC buffer by gentle centrifugation at 5,000 × g for 5 minutes at 4°C. Protoplasts were resuspended in 500 μL SuTC buffer and counted using a haemocytometer, then diluted to 10^7^ protoplasts per mL for transformation experiments.

Cell wall regeneration monitoring was performed using calcofluor white staining at 0, 1, 2, 3, and 4 hours post-isolation to assess cell wall reconstruction kinetics and determine optimal transformation timing.

### Fungal DNA extraction and target gene cloning

Approximately 100 mg of fresh mycelium from solid medium was frozen in liquid nitrogen and then homogenized using a bead mill homogenizer (Precellys 24 Touch) at 6800 rpm for 40 seconds. The DNA extraction procedure followed the instructions from the FavorPrep™ Plant Genomic DNA Extraction Kit (FAVORGEN). Quality and quantity of DNA were measured using the ratio of wavelength 260/230 and 260/280 by Nanodrop (Biometrics).

The polymerase chain reaction (PCR) was performed with 2X Taq DNA Polymerase Master Mix (AMPLIQON), and the reaction setup was according to the instructions. The PCR products were loaded in 1% agarose gel containing 0.01% safety staining dye (SYBR Safe DNA Gel Stain, Invitrogen) for running gel electrophoresis. The ideal size of bands was purified by gel purification kit (Favorgen), and the quality of purified PCR products were examined by Nanodrop.

For FoURA5 cloning, 1 µL of purified PCR product was mixed with TOPO TA Cloning Vector (Thermo Fisher) according to the manual script. After incubating at room temperature for 5 minutes, 2 µL reaction was added in 50 µL DH5_a_ competent cells (ECOS™), and the following procedure is based on the standard heat-shock transformation protocol. A primer pair for amplifying the coding region of FoURA5 was listed in Table S1.

### gRNA design *and in vitro* validation

Target site selection was performed using the *FoURA5* gene sequence as the template for CRISPR/Cpf1 targeting. The *URA5* gene encodes orotate phosphoribosyltransferase, essential for pyrimidine biosynthesis, and serves as an effective negative selection marker when disrupted. Potential guide RNA target sites were identified using Benchling CRISPR design platform (Benchling Inc., San Francisco, CA), with selection criteria including optimal TTTV protospacer adjacent motif (PAM) positioning and minimal predicted off-target effects within the *F. oxysporum* genome.

Two guide RNAs were designed: gRNA_244657 and gRNA_245247, targeting distinct sites within the FoURA5 coding sequence. Guide RNAs were synthesised as 23 nucleotide (nt) sequences with 20 nt additional scaffold specific for Cpf1. Synthetic guide RNAs were obtained from MDBio (Taichung, Taiwan) and dissolved in nuclease-free water to 100 μM stock concentrations.

*In vitro* cleavage assays were performed to validate gRNA activity before transformation experiments. Genomic DNA template was prepared by PCR amplification of the FoURA5 target region using primers P1 and P5, generating a 2,062 bp fragment containing both gRNA target sites. Cleavage reactions contained 500 ng template DNA, varying concentrations of gRNA (0.003-30 μM), and 5 μg LbCas12a protein (EnGen® Lba Cas12a, New England Biolabs M0653S) in 1× reaction buffer. Reactions were incubated at 37°C for 1 hour, then analysed by agarose gel electrophoresis to assess cleavage efficiency. Expected cleavage products were 1,203 bp for gRNA_244657 and 1,473 bp for gRNA_245247.

Concentration optimisation experiments demonstrated complete target cleavage at gRNA concentrations between 1-3 μM, with detectable activity maintained at concentrations as low as 0.15 μM. Based on these results, gRNA_244657 showed superior cleavage efficiency and was selected for subsequent transformation experiments.

### CRISPR/Cpf1 ribonucleoprotein complex assembly

Ribonucleoprotein (RNP) complex preparation was performed immediately prior to protoplast transformation to ensure optimal activity. Pre-assembly reactions contained 500 ng of selected guide RNA (gRNA_244657), 5 μg LbCas12a protein (EnGen® Lba Cas12a, New England Biolabs M0653T), and 1× reaction buffer in a total volume of 20 μL. The mixture was incubated at 25°C for 15 minutes to allow RNP complex formation, following established protocols for ribonucleoprotein assembly.

Donor DNA construct preparation involved generating EGFP reporter cassettes with varying homology arm lengths to systematically evaluate HDR efficiency. The EGFP expression cassette was amplified from pAN7.1-EGFP vector and flanked with FoURA5 homology sequences of 20 bp, 100 bp, and 1,000 bp at both 5’ and 3’ ends. For 20 bp homology arms, sequences were directly incorporated into PCR primers during amplification. For longer homology arms, sequential PCR reactions were performed using nested primer strategies to extend homologous sequences progressively.

PCR amplifications were performed using KAPA HiFi DNA Polymerase (KAPA Biosystems) according to manufacturer’s instructions, with products purified using PCR purification kits (FavorPrep, FAVORGEN). All donor DNA constructs were verified by agarose gel electrophoresis and quantified using spectrophotometric analysis before use in transformation experiments. Typical donor DNA concentrations of 200-500 ng/μL were used, with 2 μg total donor DNA per transformation reaction.

Transformation mixture preparation involved combining pre-assembled RNP complexes with 200 μL protoplast suspension (10^7^ cells/mL) and 2 μg donor DNA in a 50 mL tube. The mixture was incubated at room temperature for 20 minutes to allow RNP-protoplast interaction before PEG-mediated transformation.

### Protoplast Transformation and Selection

PEG-mediated transformation was performed using a modified protocol optimised for F. oxysporum protoplasts. Following RNP-protoplast incubation, 1 mL of PTC solution (60% polyethylene glycol 4000 in SuTC buffer) was slowly added dropwise to the transformation mixture whilst gently flicking the tube bottom to ensure gradual mixing. The transformation reaction was incubated at room temperature for 20 minutes to facilitate membrane permeabilisation and DNA uptake.

Cell recovery and regeneration involved diluting the transformation mixture with 5 mL liquid TB3 medium and incubating at 28°C for 12-18 hours to allow cell wall regeneration and metabolic recovery. This recovery period is critical for protoplast viability and subsequent selection marker expression.

Selection procedures utilised the URA5 disruption phenotype for identifying successful transformants. M100 selective medium was prepared by melting 45 mL M100 base medium to 55°C and adding 5-fluoroorotic acid (5’-FOA) to 2 mg/mL final concentration, along with 250 μM uridine and uracil as metabolic supplements. The recovered transformation mixture was gently mixed with the selective medium and poured into sterile Petri dishes before solidification. Plates were incubated at 28°C for 10-14 days to allow colony development.

EGFP-based primary screening enabled rapid identification of transformants containing integrated donor DNA. Colonies were examined under blue light excitation using a fluorescence stereomicroscope to identify EGFP-positive clones. The percentage of fluorescent colonies among total 5-FOA resistant transformants was calculated for each homology arm length condition. Representative EGFP-positive and EGFP-negative colonies were selected for molecular confirmation and preserved as glycerol stocks at -80°C.

Transformation efficiency was calculated as the number of 5-FOA resistant colonies per 10^6^ protoplasts transformed, with EGFP integration efficiency determined as the percentage of fluorescent colonies among total resistant transformants.

### Molecular Genotyping and Confirmation

Genomic DNA extraction from selected transformants was performed using the FavorPrep Genomic DNA Extraction Mini Kit (FAVORGEN) according to manufacturer’s instructions. Fungal mycelia were collected from 3-day-old liquid TB3 cultures and processed immediately to ensure high-quality DNA recovery.

PCR-based genotyping strategy utilised a comprehensive three-primer-pair system designed to distinguish different editing outcomes at the FoURA5 locus. Five primers were strategically positioned around the target site: P1 and P5 flanked the entire FoURA5 gene region, P3 and P4 were located within the EGFP donor sequence, and P1 and P2 spanned the 5’ homologous recombination junction. This primer design enabled systematic classification of editing events through differential PCR amplification patterns.

Mutation event classification was performed using three primer combinations with distinct diagnostic capabilities: P1+P5 amplification confirmed gene disruption events, P3+P4 amplification detected donor DNA integration, and P1+P2 amplification verified precise homologous recombination at the target junction. PCR reactions were performed using Blend Taq DNA Polymerase with the following cycling conditions: 95°C for 5 minutes initial denaturation, followed by 35 cycles of 95°C for 30 seconds, 58°C for 30 seconds, and 72°C for 2 minutes, with a final extension at 72°C for 10 minutes.

Editing outcome categories were defined based on PCR amplification patterns: (1) Simple Insertion (SI): successful amplification with all three primer pairs indicating precise HDR, (2) Incomplete Insertion (II): P1+P5 and P3+P4 positive but P1+P2 negative, suggesting partial donor integration, (3) InDel events: P1+P5 positive but P3+P4 and P1+P2 negative, indicating NHEJ-mediated mutations, and (4) Wild Type (WT): amplification patterns consistent with unedited sequence.

Molecular confirmation involved analysing at least 10-15 randomly selected transformants from each experimental condition. PCR products were visualised by agarose gel electrophoresis using 1% agarose gels stained with ethidium bromide, with molecular weight markers included for size verification. Representative PCR products were purified and subjected to Sanger sequencing to confirm sequence accuracy and verify precise integration events.

### Plant Pathogenicity Assays

Arabidopsis inoculation procedures were established to provide standardised conditions for evaluating *F. oxysporum* pathogenicity and validating edited strain functionality. Two-week-old *Arabidopsis thaliana* Columbia-0 seedlings grown on half-strength MS medium were carefully uprooted and root systems were gently washed with sterile distilled water to remove residual agar. Seedlings were transferred to 6-well tissue culture plates containing 3 mL liquid half-strength MS medium per well.

Fungal inoculum was prepared by harvesting conidia from 7-day-old cultures grown on half-strength PDA plates. Conidia were collected by flooding plates with sterile distilled water and gently scraping the surface with a sterile spatula. The conidial suspension was filtered through four layers of Miracloth and adjusted to 10^6^ conidia/mL using a haemocytometer. For EGFP-expressing edited strains, fluorescence was confirmed under blue light excitation before inoculation to ensure strain integrity.

The inoculation protocol involved adding 100 μL of a conidial suspension (10^5^ conidia final concentration) directly to the root zone of each seedling. Control treatments received 100 μL of sterile distilled water. Inoculated plants were maintained in the controlled environment chamber at 22°C under 16-hour photoperiod conditions for 7-14 days post-inoculation.

Disease assessment procedures included both qualitative and quantitative evaluations. Visual symptom assessment was performed daily, documenting leaf yellowing, chlorosis, stunting, and overall plant health status. Quantitative measurements included rosette diameter measurements using digital callipers, with at least 10 plants per treatment group. Disease severity scores were assigned using a standardised scale: 0: less than 30% wilt leaves; 1: 30%∼50% wilt leaves; 2: 50%∼70%; 3: over 70% wilt leaves.

Microscopic colonisation analysis was performed on EGFP-expressing strains to visualise fungal invasion and tissue colonisation patterns. At 7 days post-inoculation, infected root segments were excised and examined using fluorescence microscopy with appropriate filter sets for EGFP detection. Representative images were captured at different root locations including the stem base region, intercellular spaces, lateral root junctions, and vascular tissues to document colonisation diversity. Bright-field and fluorescence images were merged to provide tissue context for fungal localisation.

## QUANTIFICATION AND STATISTICAL ANALYSIS

Sample sizes and experimental replication varied by experimental objective, with all experiments performed in biological triplicates unless otherwise specified. In protoplast optimisation experiments, n represents individual protoplast preparation reactions, with at least 6 independent preparations per condition. All experiments in this manuscript had repeat at least three times. Each experiment has at least 5 replicate biological samples.

Statistical significance was defined as p < 0.05 for all analyses, with exact p-values reported where possible. All statistical analyses were performed using R statistical software (version 4.3.0) with appropriate packages for Student’s T testing. Graphs display individual data points alongside summary statistics to provide complete data transparency, with error bars representing SEM unless otherwise specified. No data points were excluded from analyses except where technical failures (e.g., contamination, equipment malfunction) were explicitly documented during experimental procedures.

## Notes

### Competing Interest Statement

The authors have declared no competing interest.

