## Supplementary material for "A Streamlined Protocol for Small-scale Protoplast Generation and CRISPR/Cpf1-mediated Genome Editing in *Fusarium oxysporum*": Suppliments

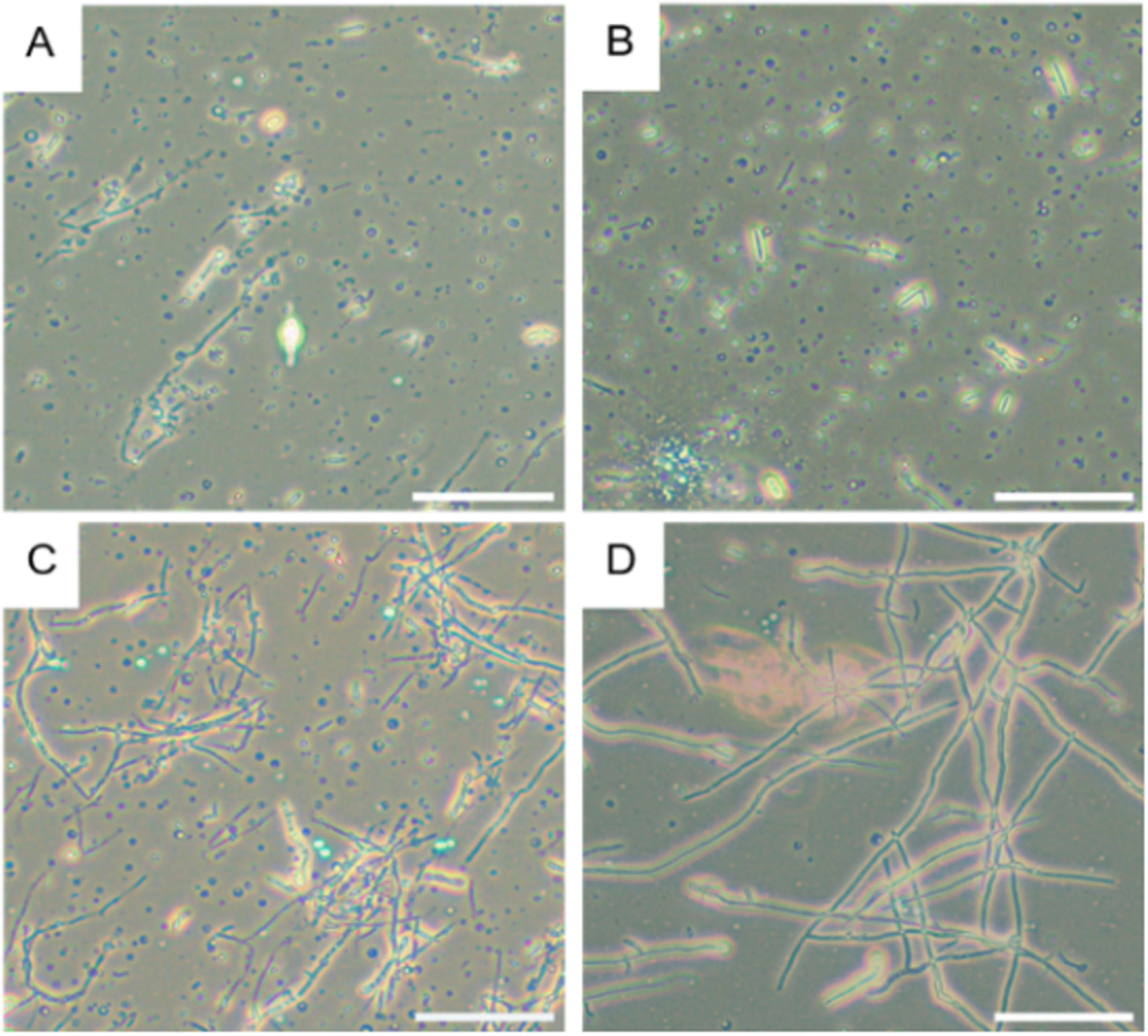


**Figure S1. Microscopic results of protoplasts in different stabliser**
Different stabliser (A: 0.8 M NaCl; B: 0.8 M KCl; C: 10 mM $Na_2PO_4$, 20 mM $CaCl_2$, 1.2 M NaCl, pH 5.8; D: 0.6 M mannitol, 10 mM $CaCl_2$, 10 mM Tris-HCl) were apply for examining the small scale protoplast generation.

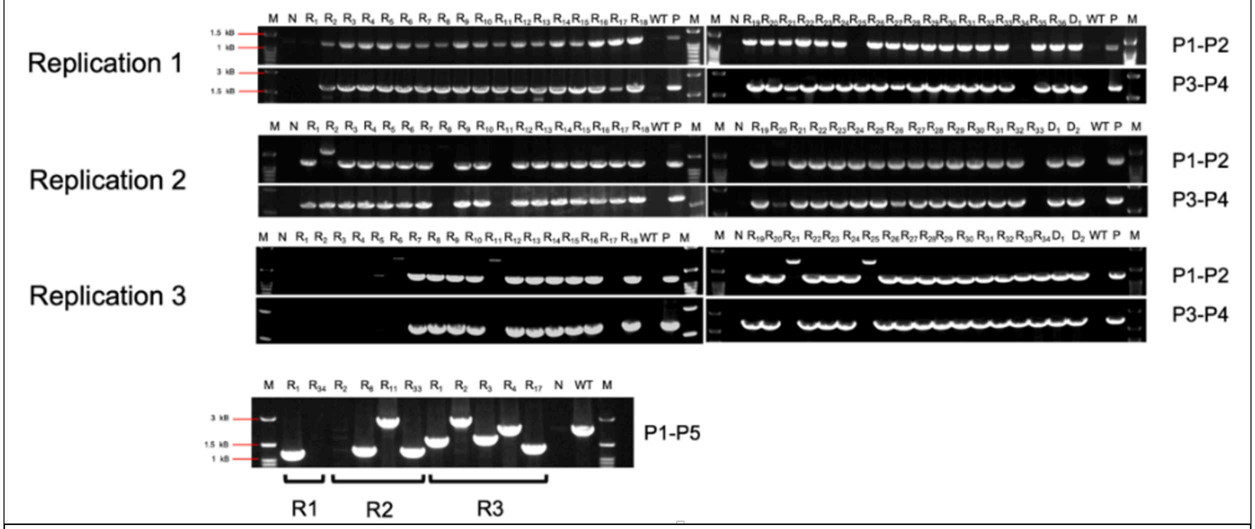


**Figure S2. Gel electrophoresis for genotyping**
Electrophoresis gel examined the polymerase chain reaction (PCR) to determine the genotyping of editing strains.

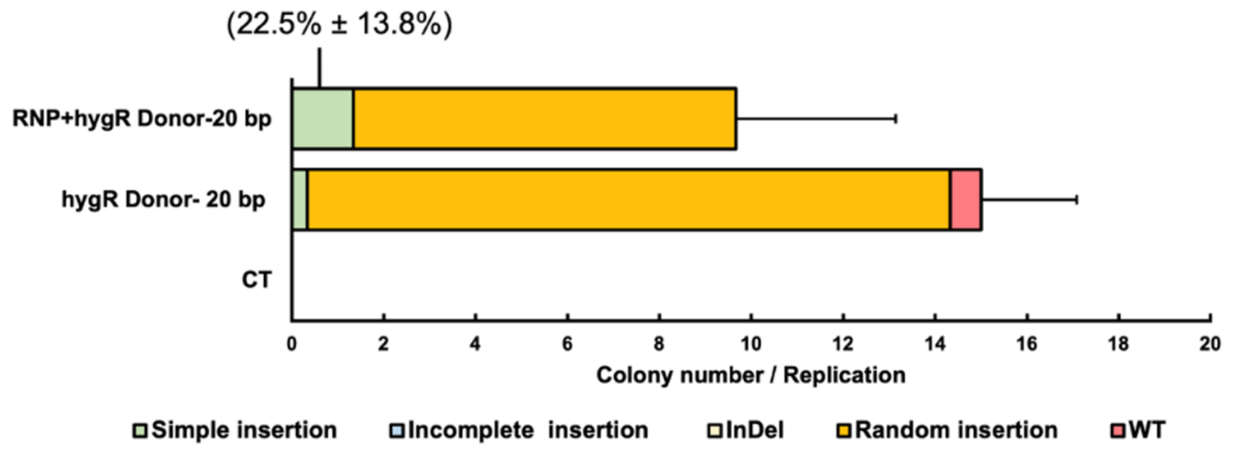


**Figure 3S. Genotyping of 20 bp homologous arms for integration the Hyg$^R$ expression donor**

The 20 bp-length homologous arms integration donor with hygromycin resistance gene, showed the variant genotypes of the edited strains.

**Table S1. Oligonucleotides use in this study**

| Name | Sequence | Source |
|---|---|---|
| gRNA_244657 | TAATTTCTACTAAGTGTAGATGAAATATTCACTCC CAATCTTGA | This study |
| gRNA_245247 | TAATTTCTACTAAGTGTAGATGCAACTCGCCAATG GCACTGGGG | This study |
| FoURA5_F | AAATGGTCGGCATCGTAGAG | This study |
| FoURA5_R | TCAAAGACCTTGGCCCAAGC | This study |
| P1 | AAATGGTCGGCATCGTAGAG | This study |
| P2 | GCTTGTGTTGTGTGACTTTTGG | This study |
| P3 | AAATATCGTGCCTCTCCTGC | This study |
| P4 | TTACTTGTACAGCTCGTCCATG | This study |
| P5 | TCAAAGACCTTGGCCCAAGC | This study |
